# Controlling Structural Bias in Intrinsically Disordered Proteins Using Solution Space Scanning

**DOI:** 10.1101/752378

**Authors:** Alex S Holehouse, Shahar Sukenik

## Abstract

Intrinsically disordered proteins or regions (IDRs) differ from their well-folded counterparts by lacking a stable tertiary state. Instead, IDRs exist in an ensemble of conformations and often possess localized, loosely held residual structure that can be a key determinant of their activity. With no extensive network of non-covalent bonds and a high propensity for exposed surface areas, the various features of an IDR’s ensemble – including local residual structure and global conformational biases – are an emergent property of both the amino acid sequence and the solution environment. Here, we attempt to understand how shifting solution conditions can alter an IDR’s ensemble. We present an efficient computational method to alter solution-protein interactions we term Solution Space (SolSpace) Scanning. SolSpace scanning uses all-atom Monte-Carlo simulations to construct ensembles under a wide range of distinct solution conditions. By tuning the interactions of specific protein moieties with the solution in a systematic manner we can both enhance and reduce local residual structure. This approach allows the ‘design’ of distinct residual structures in IDRs, offering an alternative approach to mutational studies for exploring sequence-to-ensemble relationships. Our results raise the possibility of solution-based regulation of protein functions both outside and within the dynamic solution environment of cells.

## Introduction

Nearly all biological processes occur in aqueous solutions. Yet water alone is insufficient to sustain life: in both simple and complex organisms, the composition of the surrounding solution is crucial for cellular proliferation and survival.^1^ On the molecular level, proper solution conditions are a fundamental prerequisite for protein function. Changing temperature, pressure,^2^ ionic strength,^3^ or pH^4^ can drastically alter protein structure, activity, and interactions. More nuanced changes in the makeup of the solution, such as the concentration of specific solutes, can also change protein function and stability. For example, the addition of small solutes such as urea^5^ can cause a protein to unfold and lose its native structure, even without drastically altering pH or ionic strength. In other cases, the presence of osmolytes can act as a buffer, shielding organisms from protein misfolding-related pathologies.^6^ Despite this, beyond work studying stabilizing or denaturing osmolytes, the biological repercussions of solution composition on protein structure have been largely marginalized. This stems, at least in part, from the fact that the native structure of most foldable proteins tends to be robust to changes in solution composition.^7^ Indeed, this stability is a critical reason why *in vitro* protein experiments – typically performed under conditions that are very different from the intracellular milieu – provide biologically relevant insight.

Yet not all proteins fit the same mold: intrinsically disordered proteins and protein regions (IDRs) have broken the decades-old paradigm that a well-defined structure is required for protein function.^8^ IDRs encompass an estimated 30% of the human proteome and are involved in a diverse set of cellular functions.^9^ IDRs are often described in terms of a conformational ensemble in which a collection of distinct states provides a statistical description of the intrinsic conformational biases of the sequence in question. These sequences generally contain fewer intramolecular interactions compared to folded proteins, and a majority of sidechains that are exposed to the solution.^10^ As a result, IDR behavior can be highly sensitive to changes in solution composition as intrinsic conformational biases are determined in no small part by interactions between the exposed protein surface and the solution.^11–14^

Many IDRs possess local residual structures which can alter their binding affinity or kinetics.^14–16^ We refer to ‘residual structure’ as a general term that reflects long-range sequence-specific conformational biases as well as more local interactions. Residual structure includes H-bonding and other interactions that result in local secondary structure such as helices that can facilitate structured binding interfaces, local interactions that expose or hide binding motifs, and electrostatic interactions that can upshift or downshift the intrinsic pK_a_ of charged residues. The transient nature of these interactions gives rise to a conformationally heterogeneous ensemble and introduces an inherent sensitivity to the solution environment. Taken together, this hints that even small changes to protein-solvent interactions could have a significant impact on the extent of residual structure.

Given the metastable nature of residual structure, we hypothesize that changes to the solution environment may lead to significant, sequence-specific changes to the extent and type of residual structure in an IDR’s ensemble. In effect, this would allow IDRs to function as sensors and actuators of changes in the cellular environment. To gain insight into the way IDR residual structure is affected by solution conditions we developed and deployed a computational approach that allows us to systematically vary the protein-solution interactions for all-atom simulations of IDRs. By titrating these solution conditions and measuring the types and extents of changes to residual structure we can directly interrogate how macroscopically equivalent solution conditions influence an IDR’s conformational ensemble. We find that under certain solution conditions, specific structural preferences can be enhanced, reduced, or abolished altogether, and that while global properties such as the radius of gyration or end-to-end distances may appear invariant, residual secondary structure and transient long-range interactions can be significantly altered.

The work is arranged as follows: First, we introduce the computational framework through which solution space is defined and sampled. We then demonstrate that the limited changes we make in solution space do not cause well-folded proteins to unfold. We next examine several well-studied IDRs to explore how changes to their solution environment can influence residual structure. We show that this residual structure can vary dramatically from even small perturbation in solution space. Finally, we demonstrate that the unfolded state of otherwise well folded proteins shows less variations in response to changes in solutions space when compared to the IDRs tested here, suggesting that certain IDR sequences evolved to be responsive to solution changes.

## Methods

Experimentally, the interaction between a protein and the solution around it can be quantified by the transfer free-energy (TFE):^17^ the free-energy cost associated with transferring a protein conformation from one solution to another. If, in a given solution, one conformation has a lower TFE than another, that conformation will be preferred (**Fig. 1a**). Tanford^17^, Bolen^18,19^, Record^20,21^, and others have empirically shown that, for many cases, the TFE of a protein configuration can be quantitatively predicted by measuring the solvent-exposed area of chemically distinct surface groups. The selection of surface groups can be arbitrary but is often defined by the 19 amino acid sidechains and the peptide backbone. Surface group TFE (GTFE) is experimentally accessible via solubility measurements of model compounds.^22,23^ The TFE of a protein conformation, *ΔG*_*tr*_, is thus given by

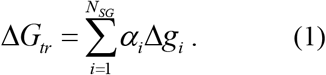

**Figure 1.**
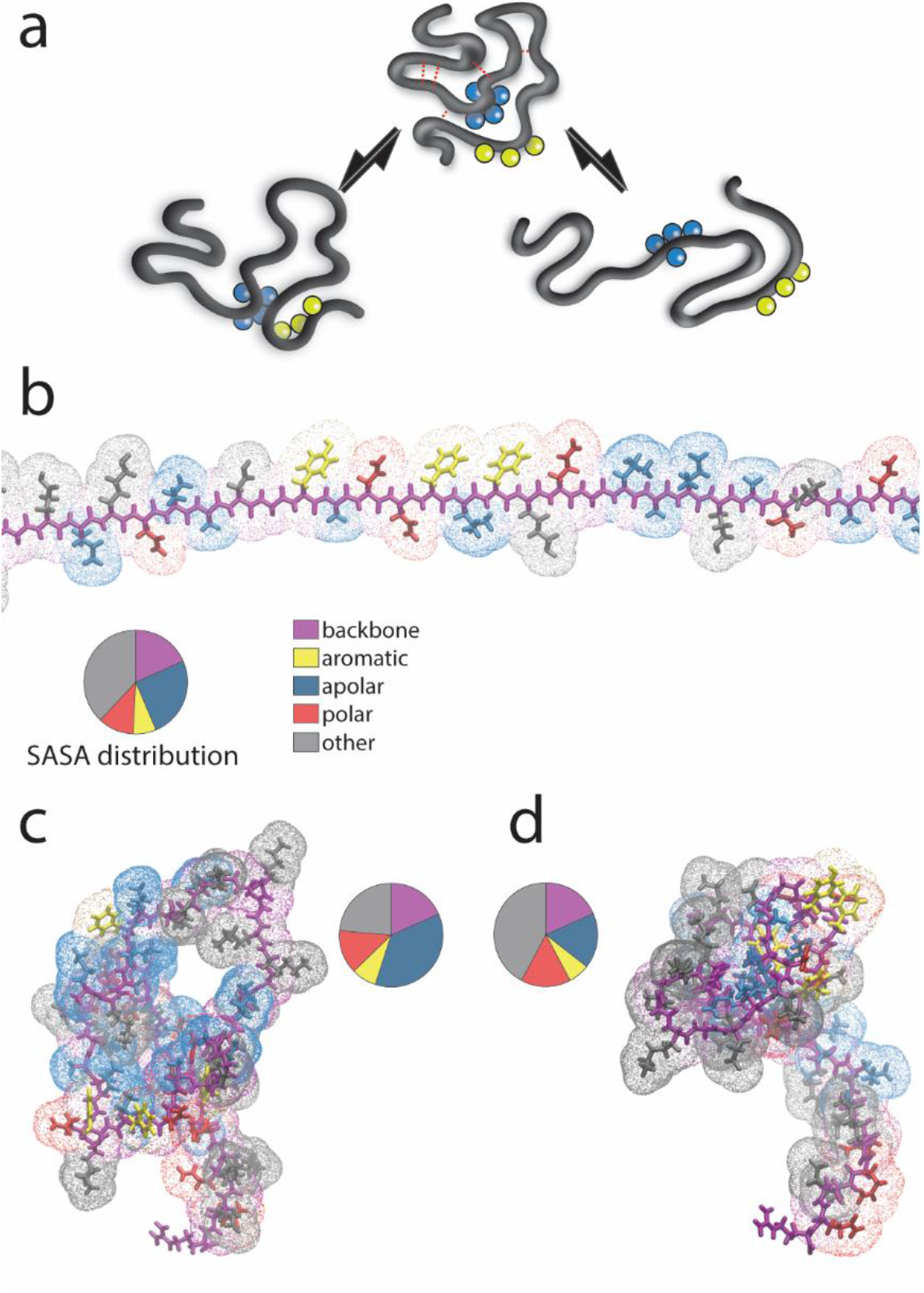
Principle of solution space scanning. (**a**) An arbitrary conformation in water hides apolar residues (blue) and exposes aromatic residues (yellow). Additional bias to this conformation is given by non-covalent bonds (red dashed lines) that promote the formation of residual structure. In a solution for which solvent-apolar residue interactions are sufficiently attractive the ensemble will be biased to expose apolar residues (right equilibrium), breaking the non-covalent bonds within the protein to form new ones with the solution. When the solution is sufficiently repulsive to aromatic residues, conformations where aromatic residues are buried is preferred (left equilibrium) (**b**) A maximally extended conformation for a given sequence is used to calculate the maximum transfer free energy 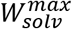. We classify each of the amino acid sidechains and the backbone into one of five possible groups (shown in the legend), as indicated by the color of the point clouds around the stick structure. The pie chart represents the different solvent-accessible surface area (SASA) fractions of these groups. **(c, d)** Representative conformations from solutions with attractive (**c**) or repulsive (**d**) interactions with apolar residues show exposure or burial of hydrophobic residues. The pie chart represents the different SASA fractions of these groups.

Here, *N*_*SG*_ is the total number of surface groups, *α*_*i*_ is the total surface area of group *i* in the conformation, and Δ*g*_*i*_ is the GTFE of group *i* per area unit. While the summation shown in Eq. 1 neglects 3-body and higher-order interactions between solution and surface types, it nonetheless manages to faithfully reproduce experimentally determined values for the free energy of folding in 2-component solutions of denaturants or osmolytes.^18,20,23,24^

We investigate the impact of solution-protein interactions on the conformational behavior of proteins by tuning GTFEs in the ABSINTH implicit solvent model. In ABSINTH, proteins-solution interactions are quantified by a mean-field implicit solvent term. Changing GTFE terms is a way to explore the space of chemically distinct solution conditions^25,26^ which we refer to herein as “solution space”. This is an analogy to the “sequence space” that is studied by mutating or shuffling the wild-type sequence of a protein.^27,28^ The ABSINTH potential energy function (which for convenience we refer to as the Hamiltonian) provides a way to calculate the instantaneous potential energy associated with a given conformation. The forcefield is used with Metropolis Monte Carlo based sampling, in which a new conformation is generated at random by perturbing some degree of freedom, and the conformations are accepted or rejected based on the difference in energy between an existing and new state and the current thermal energy. Eq. (2) defines the general form of the ABSINTH Hamiltonian:

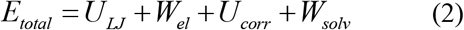

Here, the total energy of the system, *E*_*total*_, is the sum of distinct contributions: *U*_*LJ*_ represents the short range repulsive and dispersive steric interactions (Lennard-Jones interactions). *W*_*el*_ defines electrostatic interactions, which are modulated by the mean-field dielectric. *U*_*corr*_ is a torsional correction term incorporating torsion angle biases that originate from local electronic effects, such as the planarity associated with the tyrosine hydroxyl. Finally, *W*_*solv*_, is the energy of the interactions between the protein configuration and its surrounding solution. In the ABSINTH model, each polypeptide is subdivided into a list of experimentally determined GTFEs.^18^ Similar to **Eq. 1**. *W*_*solv*_ can be written as:

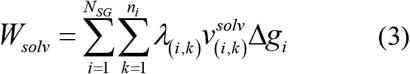

Here, *N*_*SG*_ represents the number of solvation groups in the system, *n*_*i*_ is the number of atoms in that solvation group, λ_(i,k)_ represents an atomic weighting factor for atom *k* in solvation group *i* which lies between 0 and 1 (and reflects the relative atomic radii), 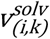 represents the solvation status (which reflects the normalized solvent accessible volume of atom *k* in solvation group *i*) and Δ*g*_*i*_ represents the GTFE for group *i* as in **Eq. 1**. Consequently, the influence of solvent on the total energy of a polypeptide can be controlled by altering the GTFE for different sets of amino acids. This provides a framework in which we can effectively modulate how individual or groups of amino acids interact with the bulk solvent while leaving others unchanged. Accordingly, we refer to solvation parameter sets in which a unique combination of GTFE values are employed as representing a specific solution composition.

We call this approach Solution Space (SolSpace) Scanning. To help subdivide solution space into a computationally tractable set of distinct chemical characteristics, we have focused on four chemical identities (**Fig. 1b**) that unite several groups together: (i) backbone moieties, (ii) polar (Gln, Asn, Ser, Thr, His), (iii) apolar (Ile, Leu, Val, Ala, Met), and (iv) aromatic sidechains (Tyr, Phe, Trp). Changing the GTFEs for one of these groups means we systematically alter the GTFEs of all group members by the same amount (see **supplementary information**). At this juncture, we make no attempt to identify or describe which combination of solutes could give rise to a given solution condition. We treat the solution as a mean-field potential that defines the cost of exposing or burying specific surface types.

The absolute impact of changing the GTFEs of distinct residues by a fixed amount will strongly depend on the amino acid sequence of the protein in question. As an example, for a glycine-serine linker, changing the GTFE of aromatic residues will have no impact, while even a small change in the GTFE of the serine sidechain will have a significant influence. To provide a way to place distinct solutions and sequences on an equivalent footing for easy comparison, we took to computing a normalized change in the maximum possible TFE associated with the full protein 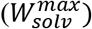.

For a given amino acid sequence this maximum possible TFE 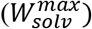 is calculated using Eq. 4 (**Fig. 1b.**)

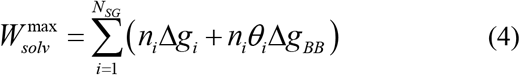

Here, *N*_*SG*_ is the number of chemically distinct GTFE groups (of which there are 20), *n*_*i*_ is the absolute number of occurrences of the *i*th group, Δg*i* is the GTFE associated with the sidechain of the *i*th group, and Δg_*BB*_ is the GTFE of the amino acid backbone and *θ*_*i*_ is a correction factor (0 ≤ *θ*_*i*_ ≤ 1) that defines the fractional solvent accessible surface area for the peptide backbone in the context of the *i*th sidechain. For example, in the case of glycine *θ*_*i*_ is 1.0 (and Δg_*Gly*_ is 0.0) while for phenylalanine *θ*_*i*_ is ∼0.6. In this way, the 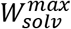 is a single value that captures the maximum possible *W*_*solv*_ (when the chain is complete extended, **Fig. 1b**) and is calculated in a manner that is agnostic regarding how those contributions are distributed across the polypeptide.

For a given sequence, the 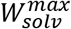 under aqueous conditions provides a useful reference point. 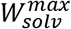 can be readily calculated under different solution conditions by varying Δg_*i*_, and this altered value can be expressed as a percentage-difference from the aqueous solution 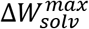,

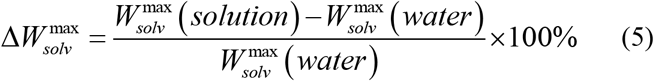

In this way, we can take two distinct sequences and for a given group of residues (e.g. apolar residues) compute the changes needed to the absolute GTFE values to provide equivalent solution for two different proteins, as assessed by 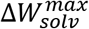. This is conceptually distinct form providing ‘the same’ solution for different proteins – instead, we are determining solutions that are energetically equivalent in terms of how they impact solvation, not in terms of molarity or composition of solutes. We use 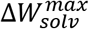 throughout the paper as a common reference point, allowing us to compare between solutions and between proteins.

For all solution space scans, we ran at least 5 independent simulations using version 2.0 of the CAMPARI Monte Carlo simulation engine over a range of distinct solution conditions (full details of simulations can be found in the Supplementary Information). The combination of CAMPARI and ABSINTH has been used extensively to characterize atomistic ensembles of unfolded and disordered proteins. In the present study, our interests lie primarily in how these ensembles change as a function of solution conditions, as opposed to the absolute ensemble behavior. Never-the-less the majority of our chosen systems (NTL9^29^, PUMA^28^, Ash1^30^) have been previously characterized extensively by ABSINTH in conjunction with Nuclear Magnetic Resonance (NMR) spectroscopy, Small Angle X-ray Scattering (SAXS), Förster Resonance Energy Transfer (FRET), and circular dichroism (CD) spectroscopy. This provides us with confidence that ABSINTH can accurately describe sequence-specific ensembles. For the systems that have not yet previously been characterized by ABSINTH we have taken advantage of extant experimental data as a touchstone for our aqueous state solution simulations. Our methodology for performing SolSpace scanning is provided as an open source Python package and the code can be accessed at https://github.com/holehouse-lab/solutionspacescanner with documentation available at https://solutionspacescanner.readthedocs.io/en/latest.

## Results

### Folded proteins do not respond strongly to changes in GTFE

How extensive is the effect of solution composition on protein structure? We first examined the native state of several well-folded single-domain proteins in different solution conditions. We scanned solution space to change the 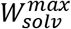 by up to ±3%, the largest change used in most of this work. This change was achieved by changing the GTFE of the backbone moiety - the major contributor in osmolytes and denaturants.^21,31^ We used these attractive or repulsive solutions to assess the sensitivity of the native structure to modest changes to solution conditions.

To assess the impact on global structure we calculated the ensemble average radius of gyration (Rg) across each solution condition, as shown in **Fig. 2a**. For perspective, a 1 M urea solution is equivalent to changing 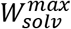 by +0.1% to +1%, depending on the amino acid composition of the protein. It is apparent that across all folded proteins and in all solutions tested we see a negligible impact on the global dimensions of the native state. We next considered the fraction of native contacts averaged across all residues (Q), and again found no significant changes across the solution conditions examined (**Fig. 2b**). Both the small change in Rg and Q indicate that folded proteins retain their native state even when the solution is strongly attractive, as shown by the high similarity to the crystal structure. While a small but noticeable change in R_g_ and Q can be seen towards 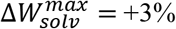, this reflects an increasingly heterogeneous folded state, as opposed to *bona fide* loss of tertiary structure (**Fig. 2c**). We further increased the 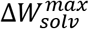 up until unfolding was observed, generating solution-dependent unfolding curves (**Fig. 2d**). These curves reveal some differences between the proteins, but the 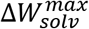 value needed to generate a 50% unfolded population was approximately between +5 and +10%. Having first established a range of 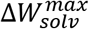 values over which folded proteins are robust, we next wondered how a set of distinct IDRs would respond over the same 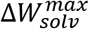 range.

**Figure 2.**
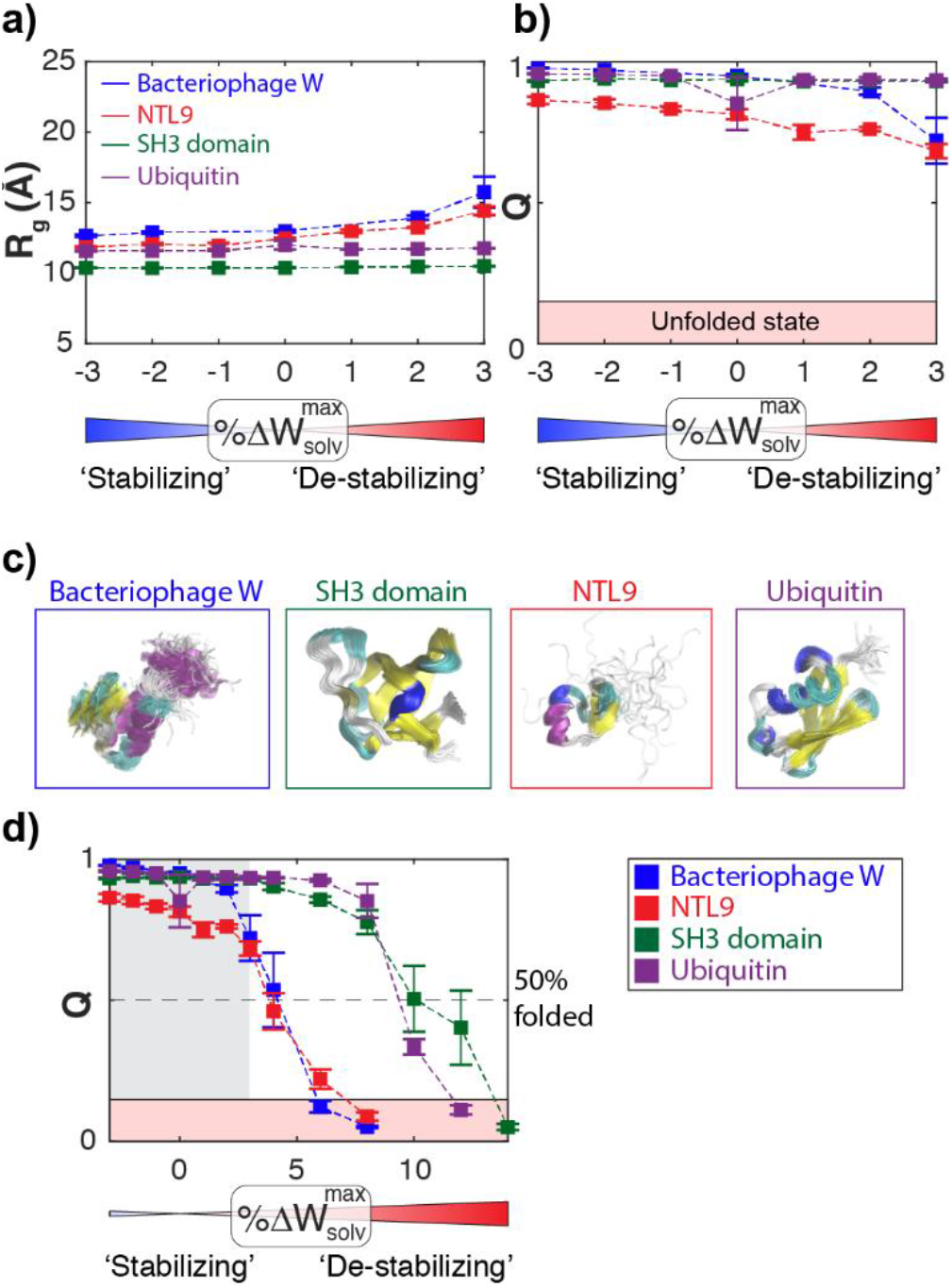
The native state is robust to solution changes. Four structurally distinct single-domain proteins were examined across a range of solution conditions. In all cases, solution is tuned by changing its interactions with the protein backbone. **(a)** The global dimensions of all four proteins are largely insensitive to changes in the solution condition, with NTL9 showing the largest variation of ∼1.5 Å. **(b)** The fraction of native contacts (Q) is also relatively insensitive to changes in solution conditions over the range explored. Note that for the unfolded state, Q is typically 0-0.1. NTL9 shows a lower Q value in general, perhaps reflecting relaxation that occurs from the starting crystal structure upon simulation in ABSINTH. **(c)** Representative snapshots from simulations with a full trajectory overlaid across the native state structure taken at 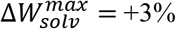. For NTL9, we see some fraying of the C-terminal helix (residues 44-56), providing a structural origin for the small increase in *R*_*g*_ with increasingly favorable 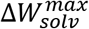. For bacteriophage W some relative motion of the local structural elements leads to a loss of native contacts, although the protein remains folded. Important, even for NTL9 the majority of the native state remains stable across 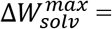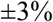 the min/max range of conditions we use throughout this study. **(d)** We further enhanced the 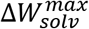 to identify the conditions under where complete unfolding is observed. For bacteriophage W and NTL9 50% unfolding was obtained at 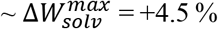, although for Ubiquitin and Src SH3 this was at 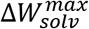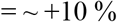. This provides some sense of the relative solution conditions that drive complete unfolding. The grey shaded area reflects the region examined in panel **b**. All error bars reflected the standard error of the mean over five or more independent simulations.

### Residual structure in intrinsically disordered proteins can be controlled by solutions

IDRs can contain residual structure – an inherent preference to adopt sequence-specific interactions that can include local secondary structure and long-range interactions.^32,33^ Altering this residual structure through mutations or changes to the solution composition has previously been shown to alter the binding affinity and kinetics of the IDR to its target.^13,34–36^ We performed extensive solution space scanning to probe how solution conditions might affect the ensemble of several IDRs whose residual structure is tied to their activity.

We first examine the N-terminal domain of p53, a protein central to the cell’s ability to prevent, correct, and respond to genomic mutations. The gene encoding for p53 is a potent oncogene and is found to be mutated in nearly all types of cancer.^37^ The N-terminal transactivation domain of p53 (p53-NTAD) comprises the first 60 residues, and has been predicted and experimentally shown to be disordered. It contains binding sites for numerous p53 activators and inhibitors, and increases the specificity of binding of p53 to its cognate DNA sequences.^38^ Borcherds *et al.* used NMR to show that p53-NTAD contains residual helical structure. The authors showed that increasing this residual helicity through mutations correlated with the increased binding affinity of the p53-NTAD to MDM2, a potent inhibitor of the full length p53 protein, both *in vitro* and *in vivo*.^34^

We wanted to see how the residual structure of p53-NTAD is affected by solution composition. To do this, we performed a SolSpace scan across a range of attractive/repulsive solutions. We were surprised to find the same 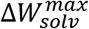 achieved using different solution interactions yielded very similar changes to R_g_, **Fig 3a**. In other words, the global dimensions of p53-NTAD are relatively insensitive to the specific details associated with the solution-protein interactions driving the conformational change. Yet the largely similar behavior in R_g_ can be misleading. To highlight this, we examined the probability to form intra-chain contacts in different solutions that give rise to the same Rg (**Fig. 3b**). The residue-residue contact maps for p53-NTAD differ depending on the identity of the solution, as shown in **Fig. 3b**. Importantly, the change in solution composition is mild 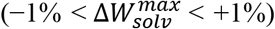. This magnitude of solution interaction had no effect at all on contacts in well folded proteins (**Fig. 2b**). Having identified difference in intramolecular contacts, we wondered how these solutions might influence sequence helicity.

**Figure 3.**
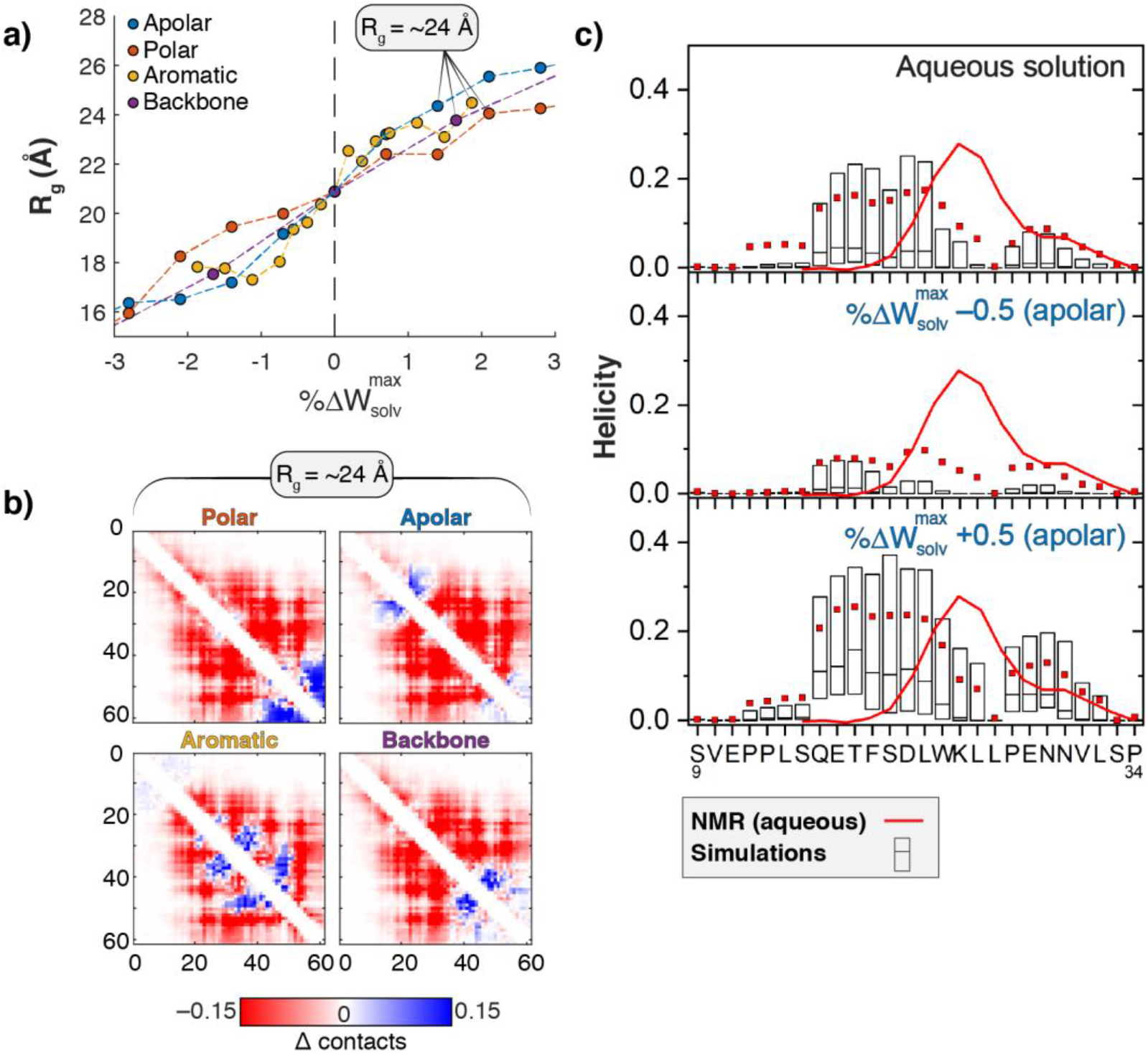
p53 N-terminal domain (p53-NTAD) residual structure perturbed by solution conditions. (**a**) Radius of gyration (R_g_) of p53-NTAD as function of solution-protein interactions. At the vertical dashed line, solution interactions mimic aqueous conditions. 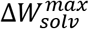 values to the right and left of this line are increasingly attractive and repulsive solutions, respectively. (**b**) Contact difference maps for different p53-NTAD ensembles at R_g_ ∼ 24 Å for different types of solutions. Flanking regions (residues 1-9 and 34-61) showed no helicity in experiments or simulations and are not shown. Contact difference maps are calculated as the as change in contact probabilities compared to aqueous conditions. Distinct contact patterns are observed across the four solution conditions despite approximately identical global dimensions. (**c**) Probability for helix formation in aqueous conditions (top) and solutions that repel 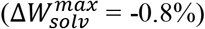 or attract 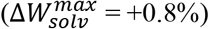 apolar residues (middle and bottom). Red squares denote averages, boxes denote 25% and 75% of the data, with the median shown as a line. N=30 independent simulations for each condition. Red line denotes the NMR-determined residual helicity of p53-NTAD reported by Borcherds *et al.*^34^

Borcherds *et al.* use NMR chemical shift data to highlight a specific region between residues 20 – 30 that has an elevated tendency to form helices. They show increasing helical content increases binding affinity to MDM2. Simulations using the ABSINTH forcefield predicts a similar extent of transient helicity in approximately the same location in p53-NTAD (**Fig. 3c**). By considering equivalent simulations performed using SolSpace scanning, we noticed that a solution with slight repulsion from apolar residues 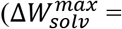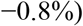 can significantly reduce the formation of helical structure, while the equivalent positive change leads to a significant enhancement in helicity (**Fig. 3c**). These results highlight the fact that small changes to solution conditions can lead to changes in residual structure to an extent that has been shown to affect MDM2 binding.

### Sensitivity to solution is encoded in IDR sequence

p53 function is in part regulated by the p53 Upregulated Modulator of Apoptosis (PUMA). The intrinsically disordered BH3 domain of PUMA (PUMA) regulates p53-mediated apoptosis by binding p53 inhibitors such as MCL1 and BCL-xL.^16,39^ Wicky *et al.* have shown that the kinetics and thermodynamics of PUMA binding to MCL1 is affected by the residual helical structure in PUMA, and that the extent of residual structure is modulated by ion-specific effects (e.g. residual structure depends on ion identity rather than ionic strength).^13^ The authors focused on specific ion effects, but their findings hint at the general phenomenon of solution-mediated regulation of PUMA structure. In aqueous solution 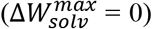, the protein displays two helical regions broken by a coil in the middle of the sequence (**Fig. 4b**), as also shown previously in experiments.^40^ Solution space scans of PUMA revealed that in aqueous conditions the protein assumes a relatively compact state, in part due to a high degree of local helicity, **Fig. 4a**. Scans also reveal that Rg is highly responsive to solution conditions. An almost complete collapse of the protein is achieved by a mild 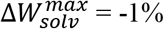 in all solution conditions tested.

**Figure 4.**
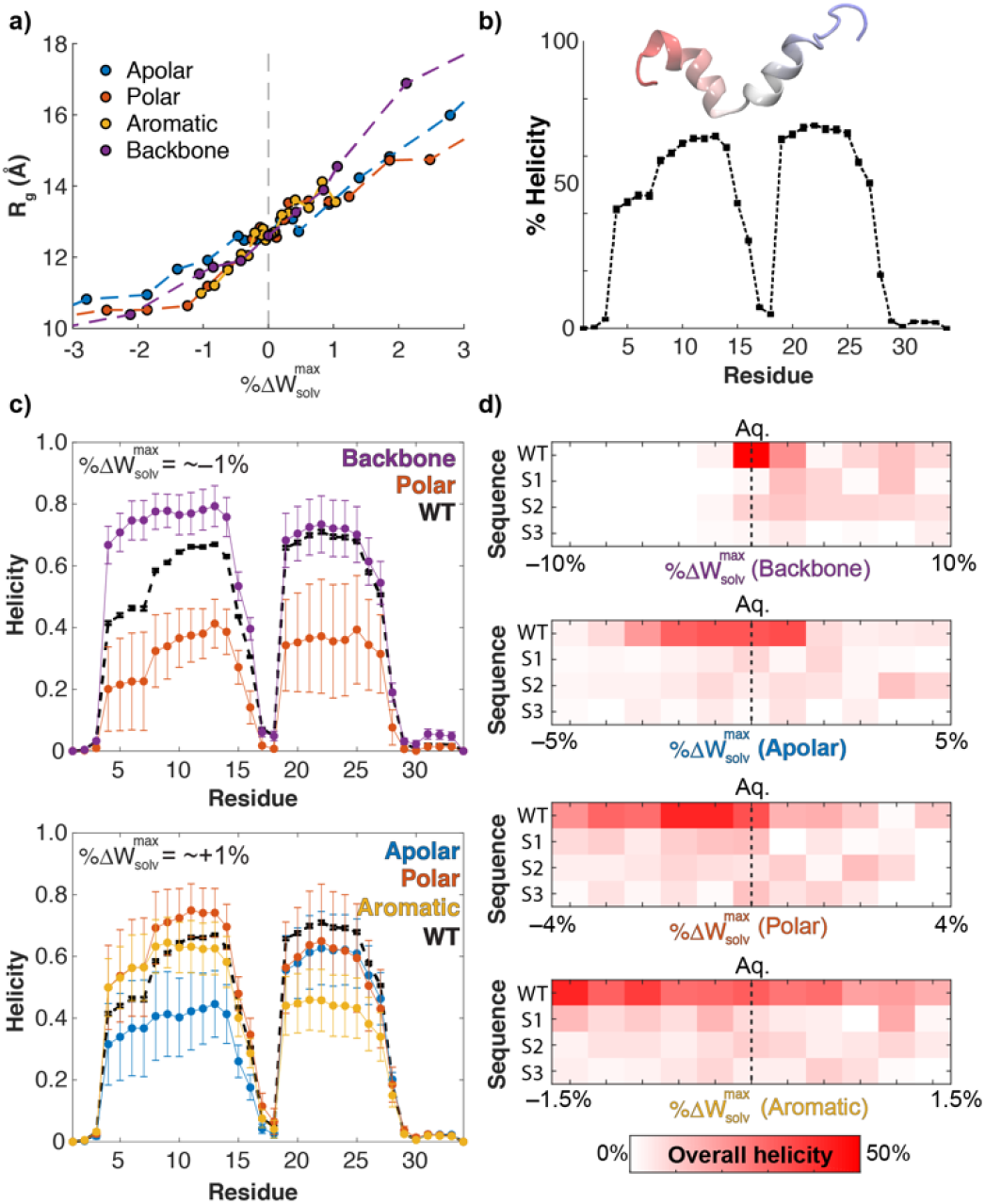
PUMA residual structure depends on its interaction with solution. (**a**) R_g_ vs 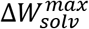 for PUMA shows a strong dependence on changes in interactions with solution. (**b**) Residual helicity for PUMA in aqueous conditions shows two helical regions divided by a short central linker. (**c**) SolSpace scan reveals solutions where the residual helicity is increased or decreased in each of the two helices, with the N-terminal helix displaying more sensitivity in both repulsive (top) and attractive (bottom) solutions compared to the C-terminal. Black symbols are same as (b) and used for reference. Error bars are standard error calculated over 5 independent simulations. (**d**) Helicity in scrambles of PUMA sequence in different solutions. Color denotes total average helicity in the sequence. Scrambles show a loss of helicity and no measurable dependence of helicity on solution conditions when compared to the WT sequence.

Examining the residual helicity in PUMA upon solution space scanning, we identified both attractive solution conditions that stabilized helicity (**Fig. 4c**, **top**) and repulsive solution conditions that reduce helicity (**Fig. 4c**, **bottom**). As before, solution interactions are changed by no more than ± 1% from aqueous solution to achieve this effect. Despite the two flanking helical regions showing equivalent extents of helicity under aqueous conditions (**Fig. 4b**), we identified several solution conditions that alter the helicity in the N-terminal half but not the C-terminal half. This correlates with Φ analysis of PUMA binding to MCL1 performed by Rogers *et al.*, which indicated that it is the N-terminal that contains more structure than the C-terminal in the transition state.^16^ These residual structure changes are significantly different from the residual structure in aqueous solution, and may alter the activity of PUMA – Wicky *et al.* show that a gain of helicity correlates with a positive change in binding affinity. Our data also shows that residual structure can increase or decrease for a similar 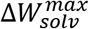 in a solution dependent manner. Thus, a −1% backbone-repulsive solution will increase N-terminal PUMA helicity by ∼20%, but the same −1% driven by polar-repulsive solutions will reduce helicity in the same region by 20% (**Fig. 4c top**).

Is the structural sensitivity of PUMA helicity encoded by the amino acid composition, or does the specific sequence in which amino acids are arranged play a role in determining local helicity and sensitivity to solutions? To answer this question, we scrambled the amino acid sequence of PUMA to generate three variants (S1-3, sequences in **Table S1**) and measured their sensitivity to changes in solution space. The *R*_*g*_ of each of the scrambles under aqueous conditions is larger than wildtype (mean scrambles = 15.1 ± 0.2 Å, mean wildtype = 12.6 ± 0.3 Å), although the response of the *R*_*g*_ to solution conditions was relatively similar across the scrambles and the wildtype (**Supplementary Fig. S1**). In contrast, for all three scrambles the residual helicity under aqueous conditions was abolished (**Fig. 4d**). Furthermore, changes in residual helicity upon solution space scanning did not occur in any sequence other than the WT (**Fig. 4d**). Taken together, our results support a general model in which the interplay between amino acid composition, specific sequence, and solution conditions dictate the extent of local helicity.

### Solutions can cause preferential burial of post-translationally modified residues

Changes to solution conditions are intimately tied to exposure or burial of specific types of surface area, as implied by **Eq. 1**.^41^ Post-translational modifications (PTMs) including phosphorylation, acetylation, methylation and ubiquitination play a wide variety of crucial roles in protein function and cellular regulation. A prerequisite for PTMs to occur is that the modification site must be exposed to the modifying enzyme.^42,43^ This is especially important for IDRs, as a large segment of post-translationally modified regions are predicted to be disorderd.^44^ While it is tempting to assume all residues within intrinsically disordered regions are equally accessible, the appearance of residual structure and the local sequence contexts means this is not necessarily the case. We wondered if changes to the solution would uniformly change the accessibility of PTM sites, or if distinct types of solution could hide some sites while exposing others.

To answer this question, we assessed the ability of solutions to modulate the exposure of phosphosites in an IDR taken from Ash1, a yeast transcription factor that controls mating type switching.^45^ The C-terminal domain of Ash1 (herein referred to as Ash1) is an 80 residue IDR that has been extensively characterized previously by NMR, SAXS and simulations under a range of solution conditions.^30^ Ash1 contains 10 phosphosites distributed approximately equally across the sequence. Upon *in vitro* incubation with a kinase nearly 100% of the sites become phosphorylated, suggesting that under aqueous solution all ten sites are sufficiently accessible for phosphorylation. This assessment is consistent with simulations performed using the ABSINTH model, which reproduce SAXS and NMR results and describes a highly expanded ensemble.^30^

To test how solution interactions alter the accessibility of Ash1 phosphosites, we performed a SolSpace scan on the Ash1 sequence. Our data corroborates the weak sensitivity of Ash1 global dimensions to denaturation – for all attractive solutions, the change in *R*_*g*_ is relatively small when compared to that of other IDRs tested. This is evident by the fast approach to a plateau in **Fig. 5a**(compare with e.g. **Figs. 4a**, **3a**). A more dramatic change is observed upon turning the solution more repulsive. In this case, Rg decreases more substantially, regardless of the driving interaction as shown in **Fig. 5a**.

**Figure 5.**
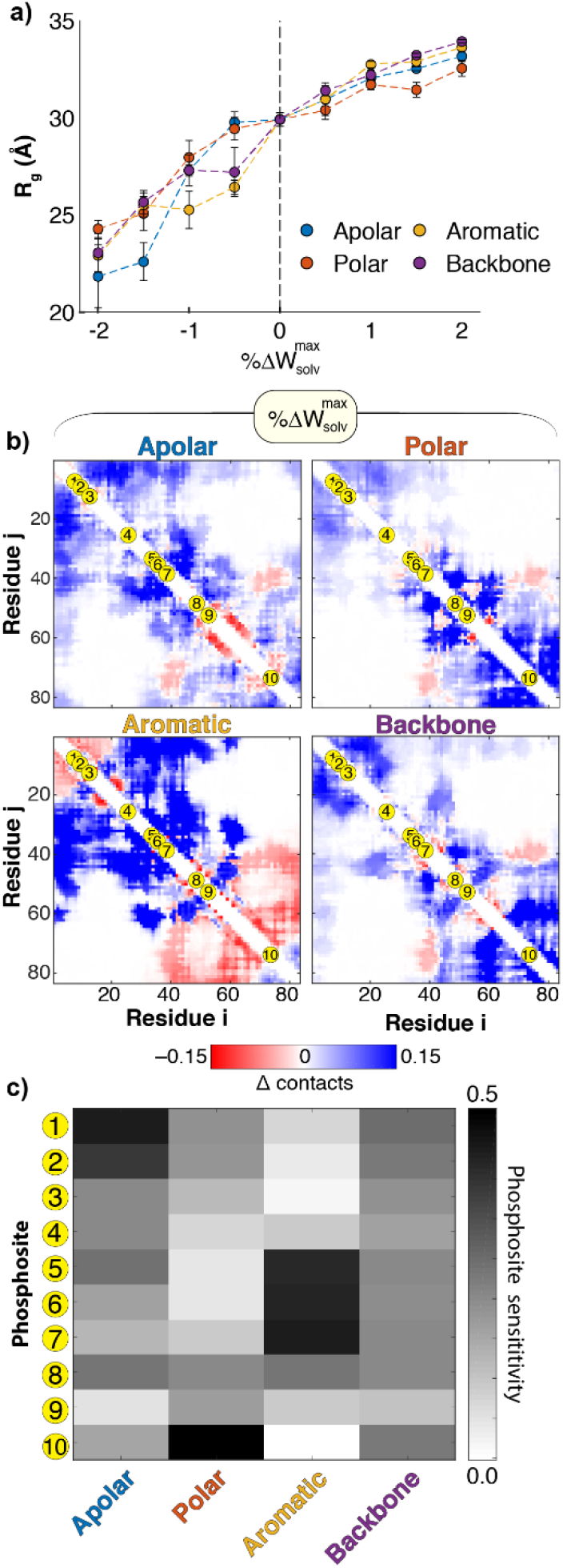
Ash1 phosphosites change their exposure in different solutions. (**a**) R_g_ vs 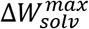 for Ash1. (**b**) Contact difference maps for different Ash1 ensembles at 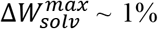 for different types of solutions. Contact difference maps are calculated as the as change in contact probabilities compared to aqueous conditions. Yellow markers on the diagonal indicate the location of the 10 phosphosites. (**c**) Phosphosite sensitivity under different solution conditions, calculated according to Eq. 6. Numbering on the left indicates phosphosite as shown in (b). Phosphosites change their sensitive (as determined by their surface exposure) depending on solution conditions.

To quantify residual structure, we analyzed contact difference maps obtained under distinct weakly repulsive solutions 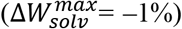. This analysis revealed distinct conformational preferences obtained under different types of solutions (**Fig. 5b**). We noticed there were specific regions that were coincident with clusters of phosphosites, hinting that certain sites might become more or less accessible depending on solution conditions. For example, under aromatic repulsive conditions, we noticed a large uptick in contacts around the 5^th^, 6^th^, and 7^th^ phosphosites as shown in **Fig 5b**, bottom left panel.

We next analyze how solvent accessibility of the 10 Ash1 phosphosites varies in different solutions. Specifically, we assessed the local region (within 3 residues) of each phosphosite *i* to calculate a fractional sensitivity (*S*_*i*_). Sensitivity was defined in terms of how the solvent accessibility under strongly attractive solution conditions 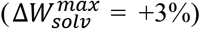 compares to the accessibility under strongly repulsive solution conditions 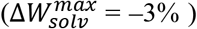, using Equation 5. The SASA used to calculate the fractional sensitivity of each site was calculated as the ensemble average SASA across the seven site residues (i−3 to i+3). A sensitivity score was calculated for each of the ten phosphosites under each of the four solution conditions.

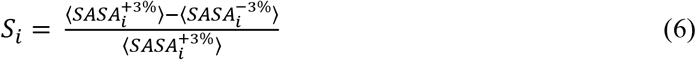

For solution conditions that vary the backbone-solvent interaction we observed uniform changes across the ten phosphosites (**Fig.5c**). This is a useful control, implying that if we titrate the backbone solubility no one region is significantly perturbed compared to another in what is intrinsically an already expanded IDR. In contrast, we identified distinct clusters of phosphosites that showed enhanced sensitivity under different solution conditions. For example, the 5^th^, 6^th^ and 7^th^ sites were highly sensitive to solutions that influence aromatic residues, but largely insensitive to those that influence polar residues. These results demonstrate that distinct solution conditions can differentially alter the solvent accessibility of specific local regions, a behavior that emerges through the interplay between chain-chain and chain-solvent interactions.

### Native-state contacts emerge under repulsive solutions of unfolded state ensembles of foldable proteins

Our analysis thus far has focused on well-characterized IDRs. Foldable proteins can also exist as unfolded ensembles under denaturing conditions and during the early stages of protein folding, and much recent attention has been focused on understanding the conformational biases within these unfolded ensembles under native conditions.^46–48^ We wondered if the unfolded state of foldable proteins under native conditions (herein referred to as the unfolded state ensemble) would show a sensitivity to solution changes similar to IDRs. We considered two model proteins that have been studied extensively in the context of protein (un)folding: NTL9 and ubiquitin.

We took advantage of recent work studying the unfolded state of the model two-state folding protein NTL9 under native conditions, for which ensembles generated by ABSINTH have been benchmarked against SAXS and FRET under native conditions.^49^ We also used ABSINTH simulations to generated a *de novo* unfolded state ensemble of ubiquitin which shows good agreement with previously published SAXS, NMR and simulation data for the unfolded state under native conditions (**Fig. S2**).^46,50,51^ We then performed a SolSpace scan on these two proteins to assess the sensitivity of the unfolded state ensembles to solution conditions.

We first assessed the global dimensions as a function of solution conditions and found results that were analogous to the IDRs we had studied (**Fig 6a**). These results suggest the global dimensions of both IDRs and the unfolded state ensemble are equivalently sensitive to solution conditions (**Fig. S4**). However, to our surprise, we observed less variation between different repulsive solutions (i.e. 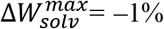) when we assessed contacts difference maps (compare **Fig. 6c, d** with **Fig. 3b** and **Fig. 5b**. The difference in variations was quantified by a matrix similarity analysis shown in **Fig. S2**, which consistently shows a higher similarity between unfolded state ensembles compared to IDR ensembles. This self-similarity in unfolded states is particularly clear across the four NTL9 ensembles, for which we are most confident in the aqueous solution state (**Fig. 6d**).

**Figure 6.**
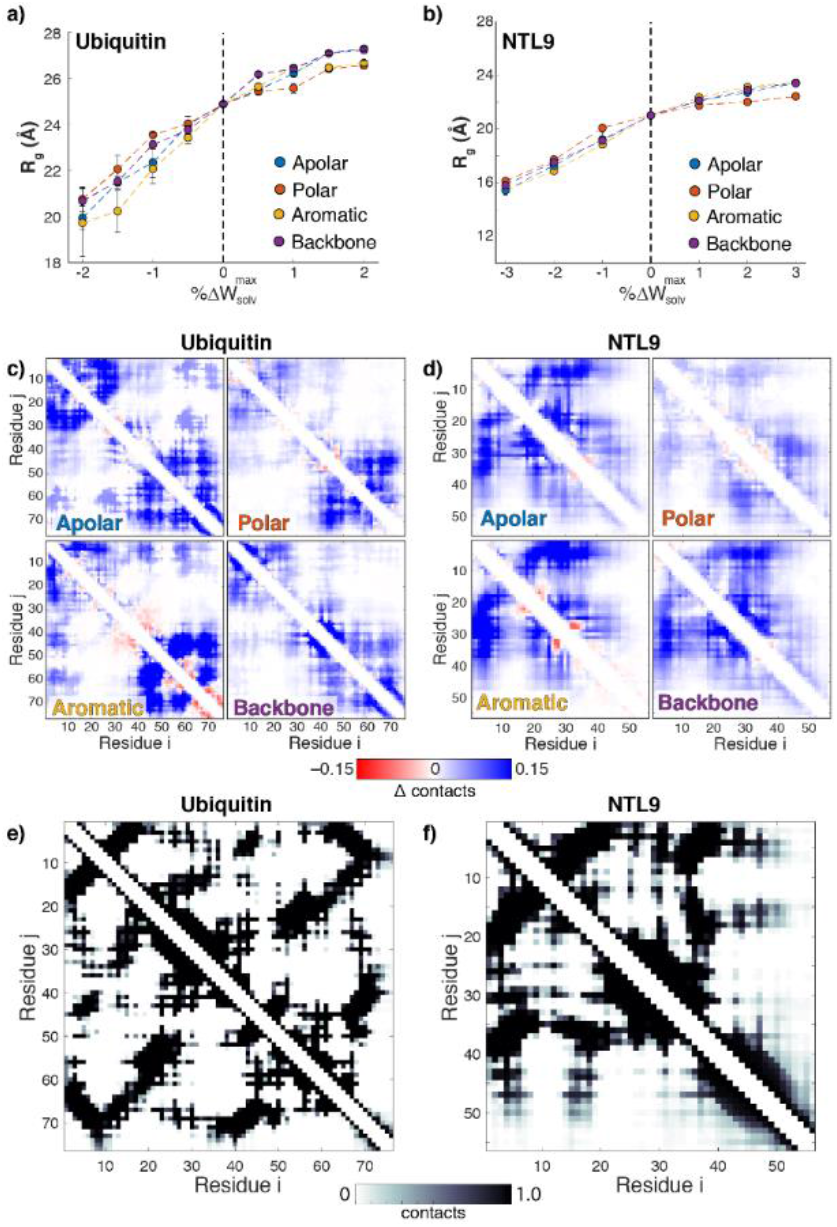
SolSpace scan of unfolded state ensembles. (**a,b**) R_g_ vs. 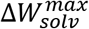 for ubiquitin (a) and NTL9 (b). (**c,d**) Contact maps for unfolded state ensembles in slightly repulsive solutions 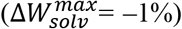 driven by different interactions for ubiquitin (c) and NTL9 (d). Solution types are noted on the bottom left hand side for each panel. (**e,f**) Contact maps for the folded state ensemble of ubiquitin (e) and NTL9 (f).

Why might chemically diverse solution environments give rise to similar contact profiles? Given folded proteins have an inherent native state, we wondered how these contacts difference profiles compared to contacts maps generated from simulations of the native state. Upon calculation of native-state contact maps (**Fig. 6e, f**) we observed a striking similarity between regions of local contacts in the native (folded) state and regions that acquire contacts under different repulsive conditions. These results suggest that the intrinsic sequence-encoded energy landscape of folded proteins is sufficiently strong such that even as solution conditions become unfavorable in different ways a ‘downhill’ route to the folded state represents the most energetically favorable ensemble. In much the same way that denaturants act as sequence-independent osmolytes that weaken the native state, a range of chemically diverse stabilizing osmolytes would be expected to strengthen the native state, a result borne out by a numerous studies showing how entirely unrelated osmolytes, proteins, and polymers can help stabilize the native state despite a wide range of underlying chemical interactions.^52–54^

## Discussion

Solution Space Scanning provides an intuitive, easy to use, and accessible method to study the sensitivity of disordered sequences to solution changes. It can be applied to any sequence and requires relatively modest computational resources to run. The predictions obtained by our method can be readily tested experimentally: The GTFEs in single-solute solutions is known for certain solutes including common osmolytes and denaturants.^18^ When it is not, quantifying GTFEs of arbitrary solutions can be done using solubility measurements of model compounds.^23^

Solution effects have been known for decades to alter protein stability and kinetics. Tanford was first to take the idea of solute-amino acid interactions as measured by TFEs and use them to explain the role of solutes in protein denaturation.^55^ This idea was expanded by Bolen who used TFEs to quantitatively predict the effect of a range of non-charged solutes on the stability of a range of well-folded proteins.^18,19,56^ Record calculated GTFEs for ions and electrolytes to explain the effects of the Hoffmeister series salts on proteins and nucleic acids, quantitatively reproducing experimental results.^20,57–59^ TFEs were utilized for implicit solvation models by Karplus^60^ and in numerous modelling programs including Rosetta^61^. TFEs were also varied in simulations by Thirumalai^41,62^ and others to account for the effects of solute addition to protein stability, notably in the Molecular Transfer Model which has provided significant mechanistic insight into protein folding.^50,63,64^

TFEs are generally thought to be subtle at physiologically relevant concentrations. Indeed, to have a measurable effect for non-specific solutes (those that act by virtue of their repulsion from protein surfaces or do not bind in specific binding sites) requires concentrations of hundreds of mM. Yet often such experiments were performed on well-folded proteins for which the majority of residues are shielded from the solution. The intramolecular bonds of folded proteins are scarcely affected by the relatively small forces of solution interactions, a result that helps rationalize why folded proteins are stable across an incredibly wide range of physiological and non-physiological solution conditions. In keeping with this assessment, our solution space scans showed little or no effect on small well-folded proteins, even at 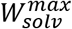 changes that would have an unfolded protein completely extended (**Fig. 2**).

The sensitivity of IDRs to solution composition has prompted many experimental studies attempting to measure the effect of changing solution conditions. Many of these focused on the effects of crowding^65–67^ – the entropically-driven force driving polymers to compact when placed in an environment with limited free volume.^68,69^ This type of entropic crowding can only act, as far as theory predicts, to compact chains. Experimentally, the picture that emerges is far more complex, and IDRs were shown to compact,^12,70^ show little change in global dimensions,^11,71^ or even expand upon exposure to crowding conditions.^72^ This complex spectrum of behavior points to the fact that steric repulsion is not adequate on its own to describe solution composition effects on heteropolymers such as IDRs even *in vitro*, let alone the complex and heterogenous cellular environment. Solution space scanning highlights the wide range of IDR responses to such non-steric, soft interactions.^73–75^ Our results also help demonstrate why physics-based models will continue to play a key role in understanding how IDRs interact with their environment; the impact of changes to the solution conditions leads to both new protein-solvent interactions, but also to changes to intramolecular protein-protein interactions, effects that may be entirely masked when making observations in a single aqueous condition.

Our results highlight the power of solutions in tuning not only the global dimensions but also the residual structure of marginally stable proteins. This suggests that residual structure can be controlled not only through changes to the primary sequence (via mutations^34,76^ and post-translational modifications^77^), but also by changing solutions conditions. This means, for example, that IDRs thought to have one specific set of conformational biases in one solution may have very different biases in another, as found for numerous so-called metamorphic proteins.^78–80^ As a result, evolution may take advantage of the ability of marginally stable proteins to illicit a rapid structural response to sudden changes in environmental conditions.

In comparison to IDRs, for which we identified a variety of solution-dependent conformational biases, the residual structures in the unfolded state of NTL9 and ubiquitin were more robust to distinct solution conditions (**Fig. 6**). Interestingly, the pattern of intramolecular interactions identified in both cases showed substantial overlap with the intramolecular contacts made in the folded state. The presence of residual native structure in the unfolded state of NTL9 under folding conditions has recently been reported^49^ while native and non-native structure has been previously shown in the unfolded state of ubiquitin.^46,81,82^ In both cases, these results were interpreted to reflect a biasing of the unfolded state to aid in the folding process^83^, and are consistent with a model for protein folding in which native state interactions exert an influence even in the unfolded state.^84^ Our data shows that not only is the unfolded state biased towards native-like contacts and/or native-like topological arrangements, but that this bias is robust to changes in solution composition. This provides some explanation for the fidelity with which protein folding can occur across a range of solution conditions. If the unfolded state was significantly perturbed under non-native solution conditions, we might expect significantly variability in folding rates and major changes in the stability of protein under distinct solution conditions due relative changes to the stability of the unfolded state (regardless of how solution conditions influence the folded state directly).

We suggest that our results offer an additional putative function for IDRs, as sensors and actuators of cellular state through changes to solution conditions. In agreement with this hypothesis, IDRs have been shown to act as sensors of pH and temperature.^85,86^ Our results suggest that, in addition to these major perturbations even subtle changes to the composition of the cellular interior may be sufficient to significantly alter the conformational state of an IDR. The cellular environment can vary significantly as a function of genome-encoded differences in protein sequence and expression patterns, as well as a function of environmental differences for ‘normal’ growth conditions. A no more obvious place in which cellular conditions vary is in the context of extremophilic organisms, including thermophiles, halophiles, acidophiles and psychrophiles. Upon comparison of folded proteins from organisms that exist under extreme environmental conditions, well-defined sequence changes are often identified.^87^ Given IDRs often undergo more rapid sequence changes across evolution, it seems plausible they may provide a mechanism to help tune cellular function under varying environmental conditions by facilitating rapid genetic adaptation. Such adaptation has already been proposed in the context of IDRs in the stress response,^86,88^ but could also play an equivalent role in mediating adaptation for other types of environmental conditions such as changes to small solute, including metabolite or nutrient composition, ionic strength, or intracellular pH.

## Acknowledgements

The authors gratefully acknowledge computing time on the Multi-Environment Computer for Exploration and Discovery (MERCED) cluster at UC Merced, which was funded by National Science Foundation Grant No. ACI-1429783. ASH received a postdoctoral fellowship from the Molecular Sciences Software Institute (MOLSSI) which supported this work financially and intellectually. This work was performed while ASH was a postdoctoral fellow in the laboratory of Dr. Rohit V. Pappu (RVP) at Washington University in St. Louis and supported by the Human Frontiers Science Program (grant RGP0034/2017 to RVP).

