## Supplementary Information for "Controlling Structural Bias in Intrinsically Disordered Proteins Using Solution Space Scanning"

***Approaches to alter*** $W_{solv}^{max}$***using the SolutionSpacerScanner package***

The SolutionSpaceScanner package can be obtained from

<https://github.com/holehouse-lab/solutionspacescanner>

Within the SolutionSpaceScanner Python package we have implemented three distinct approaches through which the GTFE of a given chemical group can be defined.

*Mode 1: Δ* $W_{solv}^{max}$

For the study presented here we calculate changes to relevant groups as Δ $W_{solv}^{max}$ (the change in percentage relative to the maximum possible transfer free energy under aqueous conditions). As a reminder, the $W_{solv}^{max}$defines the summed contribution to the solvation term in the ABSINTH Hamiltonian assuming all chemical groups are fully exposed in the context of the peptide backbone. By defining an Δ%$W_{solv}^{max}$for a new solution condition, we determine the extent to which the chemical groups influenced by this solution condition would have to change such that a newly calculated $W_{solv}^{max}$would equal some specified + or – percentage value. In effect this is solving an inverse problem: “*By what amount must we change the transfer free energies for the chemical groups (i.e. amino acid sidechains or the backbone) associated with a solution group (e.g. apolar, polar, aromatic, or backbone) to obtain a given Δ* $W_{solv}^{max}$”.

For a given protein sequence the $W_{solv}^{max}$ value is inherently weighted by the frequency of each amino acid. However, the absolute offset applied to each chemical group associated with an amino acid within a given solution group is always the same. As a tangible example, should we wish to create a solution that was repulsive for apolar residues, we would identify the value (*x*) which must be added to the transfer free energy for each of the apolar amino acids sidechain chemical groups. The value *x* is uniformly applied to all chemical groups, regardless of the frequency. In this way, we ensure we are defining a solution that effects all amino acids of a given group (e.g. apolar) equivalently, taking in to account the fact that some residues will be more frequent than others in a given sequence.

We define $W_{solv}^{max}$ in the main text as shown below in equation S1:

$MTFE=\sum_{i=1}^{N_{AA}} \left( n_{i}{\Delta g}_{i}+ n_{i}{\theta_{i}\Delta g}_{BB} \right)$	(S1)

Further, we define the Δ $W_{solv}^{max}$as

$\Delta W_{solv}^{max}=100\%\times\left( \frac{{W_{solv}^{max}}^{\delta}-{W_{solv}^{max}}^{Aq.}}{{W_{solv}^{max}}^{Aq.}} \right)$	(S2)

Here ${W_{solv}^{max}}^{Aq.}$^.^ is the $W_{solv}^{max}$ under aqueous solution conditions, while ${W_{solv}^{max}}^{\delta}$ is the $W_{solv}^{max}$ determined under some non-aqeous solution conditions for which a subset of chemical groups have been uniformly offset from their aqueous values. This unknown solution condition solves the expression above to give a specific Δ $W_{solv}^{max}$value. In this way, the Δ$W_{solv}^{max}$is solved as an inverse problem by identifying the offset needed on the relevant residues to obtain a specific Δ% MTFE value (where ${W_{solv}^{max}}^{Aq.}$^.^ is calculated using equation S1, and only Δg_i_ and/or Δg_BB_ are altered).

From a numerical standpoint, $W_{solv}^{max}$ values are almost always negative – the more negative the more favorable solvent-protein interactions. As a result, if ${W_{solv}^{max}}^{\delta}$ is more negative (favorable) than ${W_{solv}^{max}}^{Aq.}$^.^ then we obtain a positive Δ $W_{solv}^{max}$value. This was a decision made such that positive Δ$W_{solv}^{max}$ values reflect stronger protein-solvent interactions while negative Δ$W_{solv}^{max}$ reflect weaker protein-solvent interactions.

Finally, to generate a parameter set that matches a specific Δ$W_{solv}^{max}$the amino acid sequence for the protein in question is required (as it is needed to calculate the MTFE). As a result, that parameter set is intrinsically only valid for the *specific* protein sequence of that composition. This means that Δ$W_{solv}^{max}$parameter sets associated with a specific Δ$W_{solv}^{max}$ value (e.g. +1 %) will have a different meaning in the context of a different sequence as the offset is weighted by the sequence composition.

In the sss.py tool the description above reflects the mode flag --fos_percentage and means the --sequence flag is also required.

*Mode 2: Fixed offset*

Fixed offset is a much simpler mode in which a fixed value is applied to the group transfer free energy for one or more specific chemical moieties. Offsets are in kcal/mol are simply added/subtracted to the values provided in the ABSINTH parameter file provided. In the sss.py tool this reflects the mode flag --fos_offset.

*Mode 2: Fixed value*

Fixed value is, again, a much simpler mode in which a fixed value is applied to the group transfer free energy for one or more specific chemical moieties. Offsets are in kcal/mol are simply added/substracted to the values provided in the ABSINTH parameter file provided. In the sss.py tool this reflects the mode flag --fos_fixed

***Simulation details***

All simulations were run using CAMPARI V2 and the ABSINTH implicit solvent model.^1^ The same move-set was used across all systems for comparison, although different box sizes and salt concentrations were used to reflect previously defined systems. Across all solution space scan simulations all simulation parameters (other than the solvation free energies of the groups being scanned) are held fixed.

*Folded state simulations*

Simulations were performed at 293 K with a total of 20 x 10^6^ production steps and 5 x 10^6^ equilibration steps. The simulation droplet radius was 100 Å or larger, and five independent simulations were performed per solution condition. Simulations were performed in 20 mM NaCl. Folded state simulations were performed from starting structures obtained from the PDB (see table S1).

We include a comment of caution here regarding the interpretation of folded-state simulations. CAMPARI is a Monte Carlo simulation engine, and as a result there exists a finite probability of proposed move unfolding the folded protein. Due to the nature of Monte Carlo moves the likelihood of ‘refolding’ that region is extremely low, such that given a sufficiently long (finite-length) simulation the protein is likely to unfold. As a result, the unfolding curves obtained in **Fig. 2d** are provided to give a sense of the relative stability of equivalently sized folded proteins under similar solution conditions and simulation lengths to those used to simulate the set of unfolded proteins examined here. However, specific features associated with the curves likely reflect a convolution of finite sampling, chain length and protein stability. To assess this in a more rigorous and quantitative manner would require a much more in-depth analysis and goes beyond the scope of this study. We provide this comment simply as a note of caution should the reader wish to further interpret these results. The primary takeaway from the simulations in **Fig. 2** reflects the fact that at +/– 1-3% Δ$W_{solv}^{max}$ there is almost no difference with simulations under aqueous conditions, a result that is distinct from the results obtained for IDRs and unfolded proteins.

*p53 simulations*

Simulations were performed at 330 K with a total of 50 x 10^6^ production steps and 5 x 10^6^ equilibration steps. The simulation droplet radius was 106 Å, and five independent simulations were performed per solution condition. Each independent simulation generates 2500 conformations, such that for each solution condition we have an ensemble of 12,500 states. Simulations were performed in 20 mM NaCl.

*PUMA simulations*

Simulations were performed at 310 K with a total of 30 x 10^6^ production steps and 10 x 10^6^ equilibration steps. The simulation droplet radius was 71 Å, and between five and fifteen independent simulations were performed per solution condition. Each independent simulation generates 2400 conformations. Simulations were performed in 20 mM NaCl. PUMA scramble simulations were performed under the same conditions.

*Ash1 simulations*

Simulations were performed at 310 K with a total of 40 x 10^6^ production steps and 10 x 10^6^ equilibration steps. The simulation droplet radius was 127 Å, and five independent simulations were performed per solution condition. Each independent simulation generates 3200 conformations. Simulations were performed in 20 mM NaCl.

*Ntl9 simulations*

Simulations were performed at 375 K with a total of 40 x 10^6^ production steps and 4 x 10^6^ equilibration steps. The simulation droplet radius was 100 Å, and five independent simulations were performed per solution condition. Each independent simulation generates 3200 conformations. Simulations were performed in 15 mM NaCl.

*Ubiquitin simulations*

Simulations were performed at 365 K with a total of 40 x 10^6^ production steps and 10 x 10^6^ equilibration steps. The simulation droplet radius was 150 Å, and five independent simulations were performed per solution condition. Each independent simulation generates 3200 conformations. Simulations were performed in 15 mM NaCl.

***Simulation analysis***

The radius of gyration is calculated in the usual way as described previously.^2^ The extent of native contacts (Q) is defined as described previously.^3^ Contact maps are calculated as the fraction of simulation frames at which any heavy-atom from a pair of residues is within a distance threshold of 10 angstroms. Helicity is calculated by classifying each residue in each frame as being helical (or not) using the DSSP algorithm.

The “similarity score” is a metric for residual structure heterogeneity between different solutions conditions. It is the sum of the product of two difference contact probability matrices at the indicated solution conditions, where the sum is performed only when contact probability changes sign:

$S=\frac{1}{N}\sum s_{i,j}$	(S3)

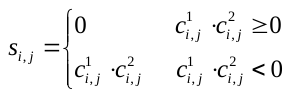
 (S4)

Here S is the similarity score, s_i,j_ is the similarity score between residues i and j, *N* is the number of residues in the sequence, and c_i,j_ is the probability difference matrix for a solution type indicated by the 1 or 2 superscript. Regions where c_i,j_ is negative indicate a distinct change in residual structure between the two solution conditions.

Synthetic scattering curves used to generate Kratky profiles in **Fig. S2** were generated using FOXS^4^ via the command:

foxs -s 400 -q 0.4 --excluded_volume 1.0 --water_layer_c2 2.0 input_ensemble.pdb -m 2

**Table S1:** Sequences used in this study (and starting PDB structures where applicable

| **Name** | **PDB ID** | **Sequence** |
| --- | --- | --- |
| **Ubiquitin** | **1D3Z** | MQIFVKTLTGKTITLEVEPSDTIENVKAKIQDKEGIPPDQQRLIFAGKQLEDGRTLSDYNIQKESTLHLVLRLRGG |
| **Ntl9** | **1DIV** | MKVIFLKDVKGKGKKGEIKNVADGYANNFLFKQGLAIEATPANLKALEAQKQKEQR |
| **Bacteriophage lambda** | **2L6Q** | MVRQEELAAARAALHDLMTGKRVATVQKDGRRVEFTATSVSDLKKYIAELEVQTGMTQRRRG |
| **SH3 domain** | **2RQT** | VRRVKTIYDCQADNDDELTFIEGEVIIVTGEEDQEWWIGHIEGQPERKGVFPVSFVHILSD |
| **p53-NTAD** | N/A | MEEPQSDPSVEPPLSQETFSDLWKLLPENNVLSPLPSQAMDDLMLSPDDIEQWFTEDPGPD |
| **PUMA** | N/A | VEEEEWAREIGAQLRRIADDLNAQYERRRQEEQH |
| **PUMA-S1** | N/A | ELARQEERGIAVHYARQEQWANQLERERDEIERD |
| **PUMA-S2** | N/A | LEWLERRRQEEVAGQEYIRDNRAEQDRIAAEEQH |
| **PUMA-S3** | N/A | IHRAIDDQYERQLERQARLVWEEEGERERNEQAA |
| **Ash1** | N/A | GASASSSPSPSTPTKSGKMRSRSSSPVRPKAYTPSPRSPNYHRFALDSPPQSPRRSSNSSITKKGSRRSSGSSPTRHTTRVCV |

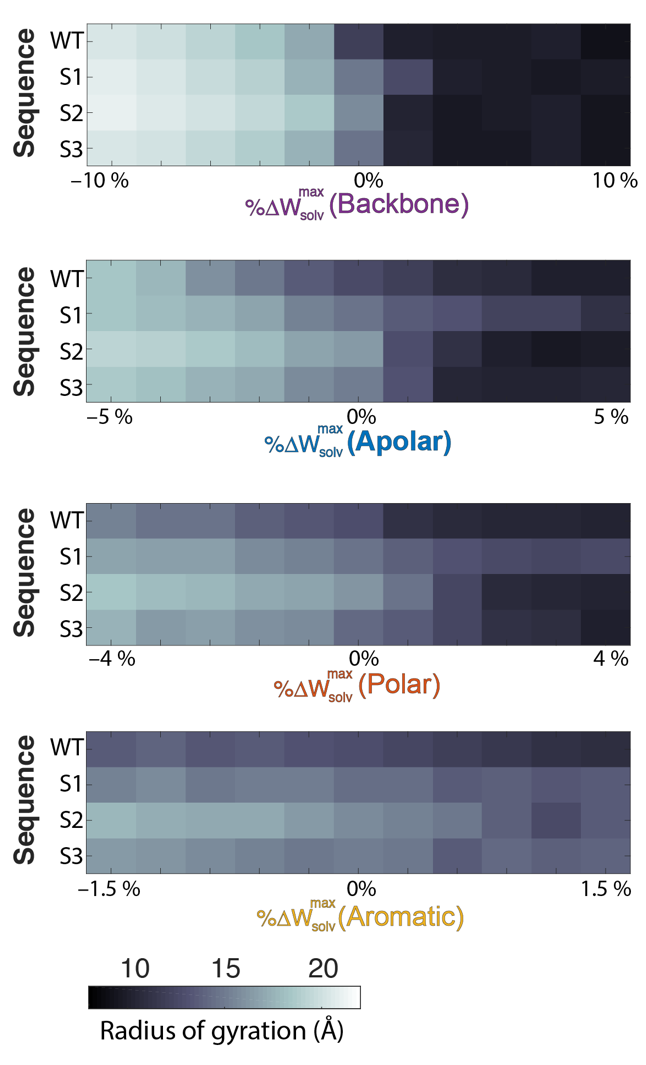

**Figure S1.** **Global dimensions of PUMA scrambles under distinct solution conditions.** Comparison of the Rg between wildtype PUMA and scrambles S1, S2, and S3 across a range of solution conditions. The wildtype sequence is generally more compact than the scrambles, owing to the higher helical content (see **Fig. 4d**). Despite this, the R_g_ is relatively consistent across the different sequences under the same solution conditions, demonstrating how equivalent global parameters are compatible with a broad range of distinct underlying conformational features.

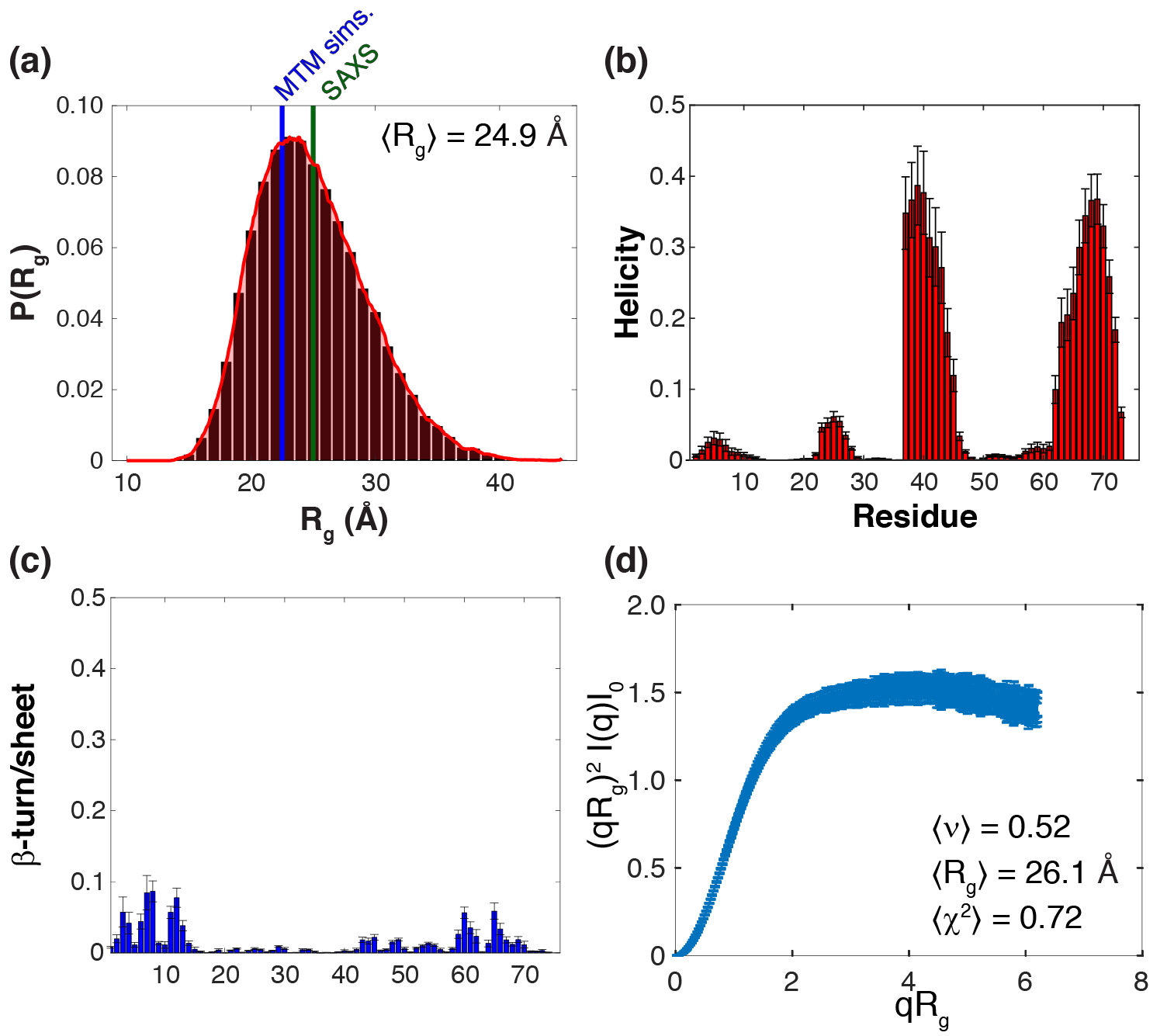

**Figure S2.** **Overview of simulation results from the unfolded state of ubiquitin.** Simulations performed with the standard ABSINTH Hamiltonian under aqueous conditions at 365K reveal an ensemble that qualitatively recapitulates a number of distinct features reported in previous studies. **(a)** Ensemble average global dimensions (24.9 Å) are consistent with those reported from time-resolved SAXS (~25 Å) and from coarse-grained simulations (22.5 Å).^5,6^ Note this smaller than unfolded state under strongly denaturing conditions, which is approximately 28 Å, consistent with a modest contraction of around 20% upon dilution into native-state conditions while simultaneously demonstrating the unfolded state is relatively expanded.^6–8^ **(b)** An analysis for residual helical structure identifies substantial non-native helicity in the C-terminus of ubiquitin under native like conditions. Residual helicity at 15-20% has been reported in the acid-denatured, methanol-denatured and urea-denatured states of ubiquitin, and our results tentatively suggest this helicity may be even more prominent in the unfolded state under native-like conditions .^9–11^ **(c)** An analysis for residual beta-turn/sheet identifies native-like beta-turn in the N-terminus, another structural feature identified in both the acid, methanol, and urea denatured states. **(d)** Synthetic scattering profiles were generated using the FOXS program and used to back-calculate dimensionless Kratky profiles. Kratky profiles show qualitative agreement with those from burst-phase SAXS measurements (a plateau at larger *q* values), measurements that describe the unfolded state under folding conditions.^5^ We also utilized a recently published molecular form factor to estimate the apparent scaling exponent (ν*^app^*) from derived scattering data.^11^ This analysis revealed an ensemble average ν*^app^* of 0.52, extremely close to values measured from smFRET (~0.50), simulations (~0.50), and to the value expected for many unfolded proteins under folding conditions (0.54).^6,9,11^ Taken together, our results suggest we have been able to generate an atomistic ensemble consistent with the unfolded state of ubiquitin under folding conditions.

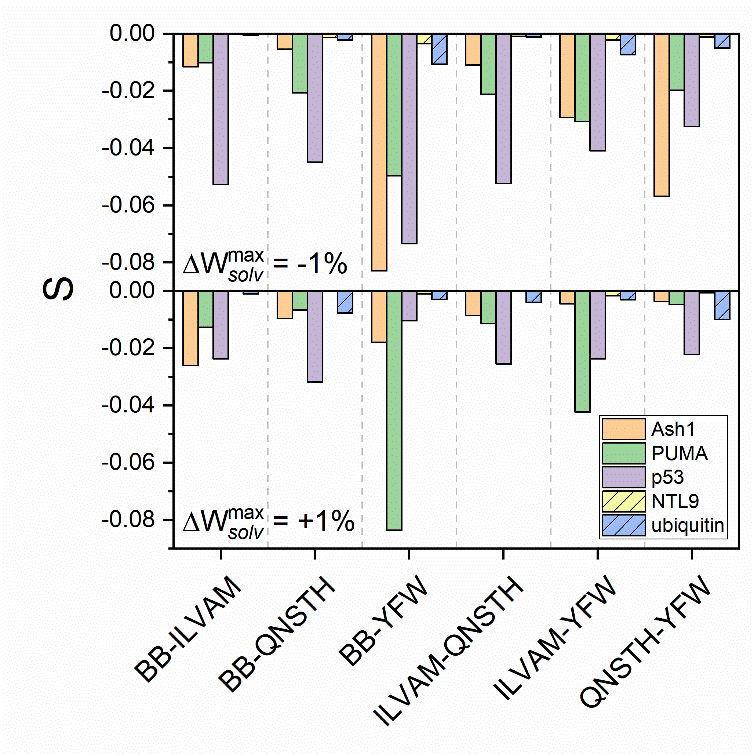

**Figure S3 –Unfolded state ensemble show more self-similarity than IDR ensembles.** The total similarity score S between different solution conditions with identical Δ$W_{solv}^{max}$ is calculated according to **Eqs. S3** and **S4**. More negative S values imply a lower similarity in the contact maps of the two ensembles specified on the abscissa. The unfolded state of NTL9 and ubiquitin shows consistently higher similarity between all solution conditions compared to the IDRs Ash1, PUMA, and p53-NTAD.

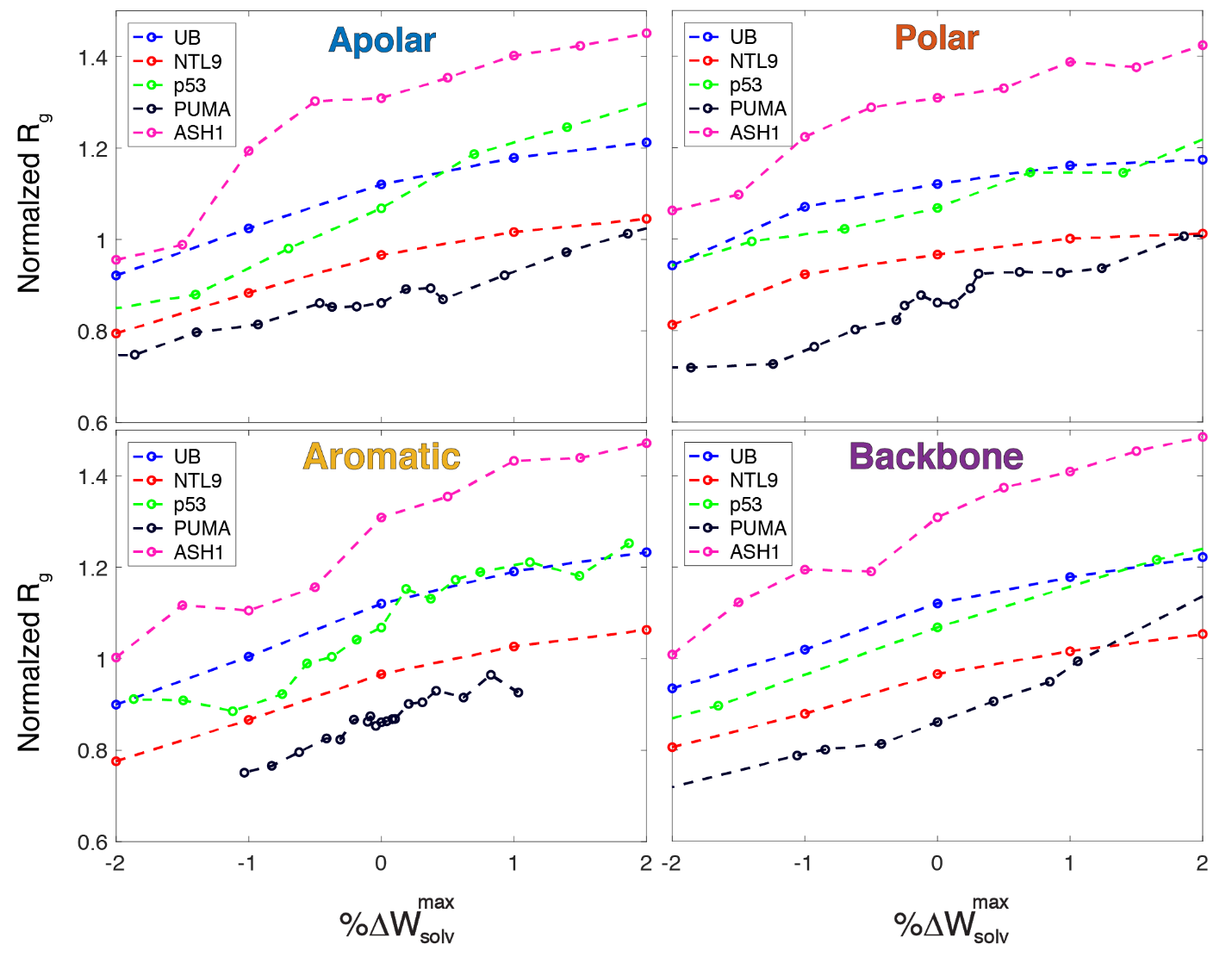

**Figure S4 – Normalized dependence of the radius of gyration on Δ**$\boldsymbol{W}_{\boldsymbol{solv}}^{\boldsymbol{max}}$**.** To allow us to compare different sequences with one another the absolute radius of gyration (*R_g_*) was normalized by the *R_g_* expected for a length and sequence-matched polypeptide simulated as a Gaussian chain (see Ref. 2 for details). For each of the five sequences examined across different conditions a relatively similar normalized *R_g_* response was observed, with sequence-specific variation as a function of solution. Importantly the unfolded ensembles of NTL9 and ubiquitin showed a global response to solution conditions that was highly analogous to the IDRs in terms of the magnitude of *R_g_* change, demonstrating that the *global* dimensions of IDRs and unfolded ensembles of normally foldable proteins show a similar response to changes in solution conditions. Despite this, as shown in **Fig. 6** and **Fig. S3**, the intramolecular interactions for NTL9 and ubiquitin are much more similar to one another across repulsive solution conditions compared to equivalent ensembles of IDRs. These results support a model in which native state interactions are present even in the unfolded state and are relatively robust to solution conditions, consistent with a folding-landscape model of protein folding.
